# Biofabrication of development-inspired scaffolds for regeneration of the annulus fibrosus macro- and microarchitecture

**DOI:** 10.1101/2021.11.03.467069

**Authors:** Nadine Kluser, Christoph Sprecher, Gion Ursin Alig, Sonja Häckel, Christoph E. Albers, Mauro Alini, Sibylle Grad, David Eglin, Andrea Vernengo

## Abstract

Annulus fibrosus (AF) tissue engineering is a promising strategy for repairing the degenerated intervertebral disc (IVD) and a research area that could benefit from improved tissue models to drive translation. AF tissue is composed of concentric layers of aligned collagen bundles arranged in an angle-ply pattern, an architecture which is challenging to recapitulate with current scaffold design strategies. In response to this need, we developed a strategy to print 3D scaffolds that induce cell and tissue organization into oriented patterns mimicking the AF. Polycaprolactone (PCL) was printed in an angle-ply macroarchitecture possessing microscale aligned topographical cues. The topography was achieved by extrusion through custom-designed printer nozzles which were either round or possessing circumferential sinusoidal peaks. Whereas the round nozzle produced extruded filaments with a slight uniaxial texture, patterned nozzles with peak heights of 60 or 120 μm produced grooves, 10.87 ± 3.09 μm or 17.77 ± 4.91 μm wide, respectively. Bone marrow derived mesenchymal stem cells (BM-MSCs) cultured on the scaffolds for four weeks exhibited similar degrees of alignment within ± 10 ° of the printing direction and upregulation of outer AF markers (COL1, COL12, SFRP, MKX, MCAM, SCX and TAGLN), with no statistically significant differences as a function of topography. Interestingly, the grooves generated by the patterned nozzles induced longitudinal end-to-end alignment of cells, capturing the arrangement of cells during fibrillogenesis. In contrast, topography produced from the round nozzle induced a continuous web of elongated cells without end-to-end alignment. Extracellular collagen I, decorin and fibromodulin were detected in patterns closely following cellular organization. Taken together, we present a single-step biofabrication strategy to induce anisotropic cellular alignments in x-, y-, and z-space, with potential application as an *in vitro* model for studying AF tissue morphogenesis and growth.

## 1. Introduction

Low back pain (LBP) is estimated to affect 568 million people worldwide (1) and is the leading global cause of disability (2, 3). There are different causes for the condition, but a strong correlation exists between LBP and the degenerated intervertebral disc (IVD) (4-6). The IVD is a composite structure consisting of a central nucleus pulposus (NP) and a peripheral annulus fibrosus (AF) (7-9). Degeneration, which is thought to originate in the NP (10), is characterized by changes in tissue matrix composition (11-14), formation of structural defects in the AF (15, 16), herniation of NP tissue (17, 18), disc height loss and biomechanical alterations (19-21). The most common surgical intervention for disc herniation, discectomy (22, 23), involves removal of extruded NP fragments to alleviate pain symptoms (24, 25). However, the surgery does not restore mechanical functionality to the IVD (26, 27) or seal AF defects (28-30), and thus herniation may reoccur (24, 31, 32). Currently, there is a clinical need for more effective methods of AF repair (30, 33).

Tissue engineering strategies comprised of cells, biomaterial scaffolds, biochemical and mechanical cues are regarded as a potential avenue for achieving AF repair. However, the highly complex, anisotropic and heirarchical structure of the native AF is a significant challenge to recapitulate with current bioengineered models. The AF is comprised of elongated, fibroblast-like cells aligned along parallel type I collagen fibers. These fibers are arranged into sheets called lamellae which wrap concentrically around the NP. The angular orientation of the collagen fibers within each lamella alternates with successive layers conferring a lattice-like, angle-ply architecture (34-36). Recent *in vitro* attempts to capture AF tissue microarchitecture have emphasized the importance of inducing elongated cell orientation within scaffolds. In multiple recent works the use of electrospun nanofibrous mats to induce cell elongation and deposition of oriented collagen fiber bundles has been studied (37-47). However, a major disadvantage hampering clinical translation of this approach is the requirement of a post-processing step, such as folding (48), stacking (21), or rolling (49) to generate 3D structures, and a tendency to exhibit limited porosity for cell viability and matrix deposition (50). Alternatively, 3D matrices composed of fibrous collagen (51-54) or fibrin (55) have been studied in combination with mechanical stretch to achieve oriented collagen matrix deposition. Overall, however, the *de novo* tissues still lack the biomechanical performance and/or nanometer-scale organization of native collagen (51, 56). Notably, current scaffold-based approaches are characterized by the random spatial arrangement of elongated cells relative to one another. This contrasts with what has been observed for embryonic fibrillogenesis, where longitudinal collagen growth occurs between neighboring cells stacked precisely end-to-end (57, 58). We suspect that the organization and mechanical functionality of engineered aligned collagen tissue may be enhanced by a new scaffold design favoring cellular organization more closely resembling developmental fibrillogenesis.

To address this need, we developed a translational biofabrication method for producing 3D scaffolds with cell instructive, microscale surface topographies that induce cellular organization into patterns mimicking fibrillogenesis. To achieve this, polycaprolactone (PCL) was printed through custom-designed 3D printer nozzles with periodic circumferential peaks. It was hypothesized that extrusion through the patterned nozzles would result in filaments posessing uniaxially aligned surface grooves inducing longitudinal cellular end-to-end alignment across the scaffolds. Furthermore, we hypothesized that layer-by-layer deposition of the grooved filaments into an angle-ply structure would enable assembly of anisotropically-oriented extracellular matrix (ECM) in a macroarchitecture resembling the native AF. Towards investigating these hypotheses, we extruded PCL scaffolds through patterned nozzles with varying sinusoidal peak heights and investigated the influence of the resulting topographies on human bone marrow derived mesenchymal stem cell (BM-MSC) alignment, differentiation and ECM expression.

## 2. Materials and Methods

All chemicals and PCL (Mw=45 kDa) were purchased from Sigma-Aldrich (St. Louis, USA) if not otherwise stated. TaqMan™ reagents and primers for real-time polymerase chain reaction (PCR) were purchased from Applied Biosystems by Thermo Fisher (Waltham, USA). Cell culture reagents alpha-minimum essential medium (α-MEM), penicillin/streptomycin (Pen/Strep), trypsin, high glucose dulbecco’s modified eagle’s medium (HG-DMEM) and non-essential amino acids were purchased from Gibco (Carlsbad, USA). Fetal bovine serum (FBS), Insulin-Transferrin-Selenium (ITS™ Premix) and ultra-low attachment 24-well plates were purchased from Corning (Corning Incorporated, New York, USA).

### 2.1 Fabrication of 3D printed constructs

3D printed constructs were fabricated by melt extrusion using a 3D Discovery printer (RegenHu Ltd., Villaz-St-Pierre, Switzerland). PCL pellets were heated to 75 °C and extruded with three different custom-manufactured nozzles: A round nozzle, with inner diameter of 300 μm, and two patterned nozzles, with inner diameters of 300 μm and periodic circumferential peaks 60 or 120 μm high, respectively (***Supplementary Figure 1***). Molten PCL was extruded through the nozzles at a printing pressure of 1 bar. PCL flowrate (revolutions per meter) was chosen proportionally to nozzle cross-sectional area with 16, 21 and 30 revs/m for the round, the nozzle with 60 and 120 μm peak height, respectively. A printing speed of 6 mm/s was used for printing the scaffolds in order to achieve high print fidelity, as higher nozzle translation speed was found to result in sagging of the struts and poor conservation of the topography (***Supplementary Figure 2***).

Scaffolds with an angle-ply lattice architecture were designed in BioCAD software (RegenHu Ltd., Villaz-St-Pierre, Switzerland, version1.1-12). The rectangular constructs (10 mm length x 5 mm width) were printed in 4 layers and with filaments oriented at angles of ± 30 °. Layer thickness depended on the nozzle used, with 0.3 mm for the round nozzle and 0.5 mm for the patterned nozzles.

### 2.2 3D Printed Scaffold Characterization

#### 2.2.1 Light Microscopy

Samples were soaked in liquid nitrogen before cutting perpendicular to the printing direction. A Zeiss light microscope (Carl Zeiss Microscopy GmbH, Jena, Germany) was used to image the cross section and top view of the 3D printed PCL constructs and peak height was measured (n ≥ 29) with AxioVision4 (version 4.9.1.0, Carl Zeiss Microscopy).

#### 2.2.2 Scanning Electron Microscopy

Scanning electron microscopy (SEM) was performed to assess scaffold surface topography. The PCL scaffolds were mounted on an aluminium stub (diameter 12.5 mm), coated with 10 nm Au/Pd (80/20 wt %) and imaged in SEM (Hitachi S-4700 II, Hitachi, Tokyo, Japan). Filament diameter (diameter of outer circumference) and groove width (n ≥ 15) were measured from SEM micrographs with ImageJ (version 1.52, National Institutes of Health, Bethesda, Maryland, USA).

#### 2.2.3 Mechanical Characterization of Printed Constructs

For mechanical characterization, scaffolds (15 × 5 mm, n=4) were 3D printed in a dog bone shape (***Figure 2(b)***) for fixation into tensile grips. A mechanical testing device (Instron 5866, Instron, Norwood, USA) equipped with a 50 N load cell was used to evaluate the tensile strength of the scaffolds. Uniaxial tension was applied to the samples at a transverse speed of 2 mm/min without preloading. Circumferential tensile modulus were calculated from the stress-strain curve in the linear region using MATLAB (Version R2020b, MathWorks, Natick, USA).

**Figure 1:**
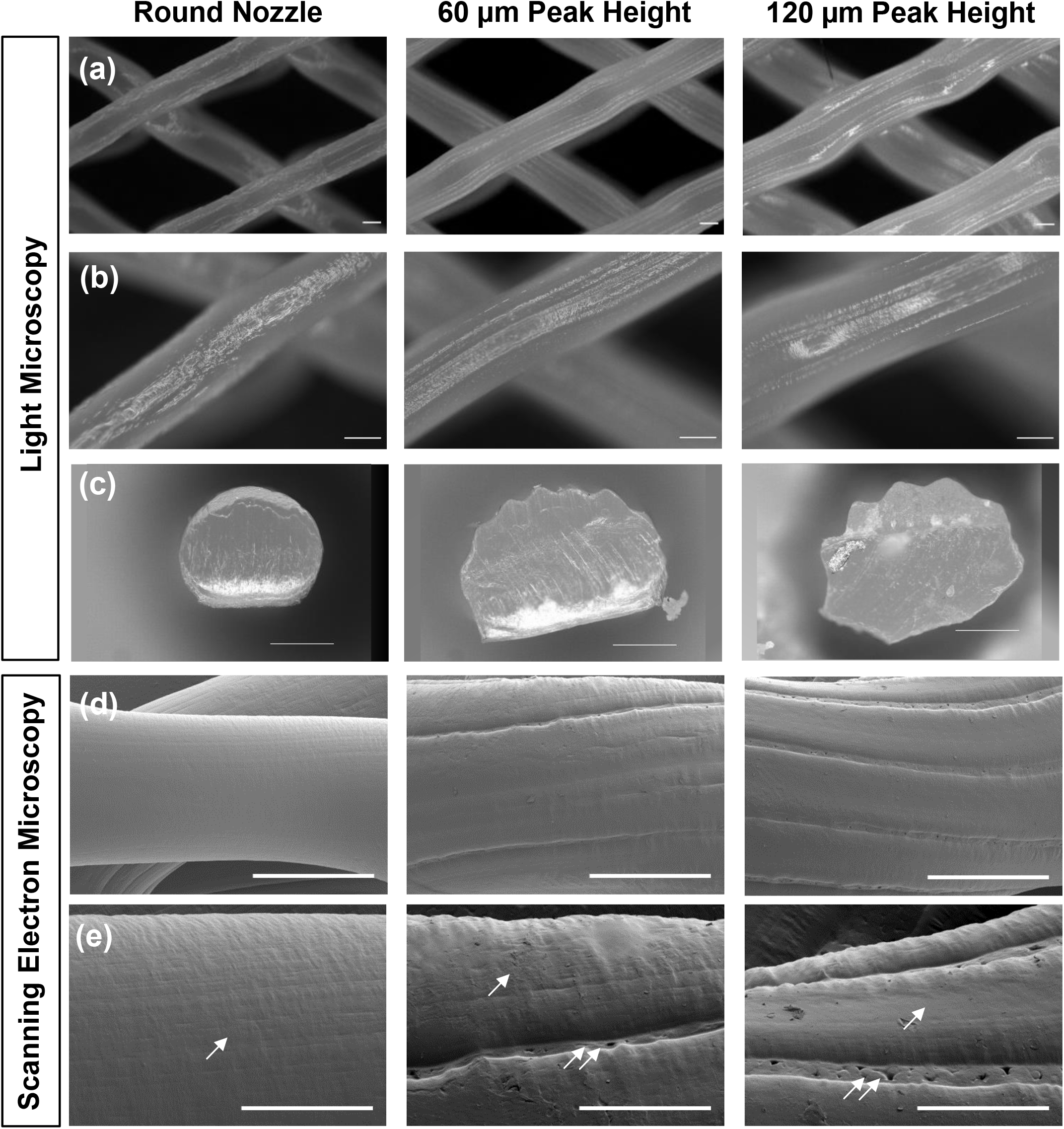
Microscopic image analysis of PCL scaffolds printed with the different nozzles (round, 60 μm or 120 μm peak height). **(a)(b)** Light microscopy top-view images of the uniaxial aligned surface topographies. (Scalebars 200 μm) **(c)** Cross sectional area of 3D printed filaments. (Scalebars 200 μm) **(d)(e)** Scanning electron microscopy top-view images of the surface topographies. (Scalebars 300 μm **(d)** and 100 μm **(e)**). Overall, all the nozzles extruded filaments possessing aligned textures (single arrows), but extrusion through the patterned nozzles additionally generated uniaxial aligned microgrooves (double arrows).

**Figure 2:**
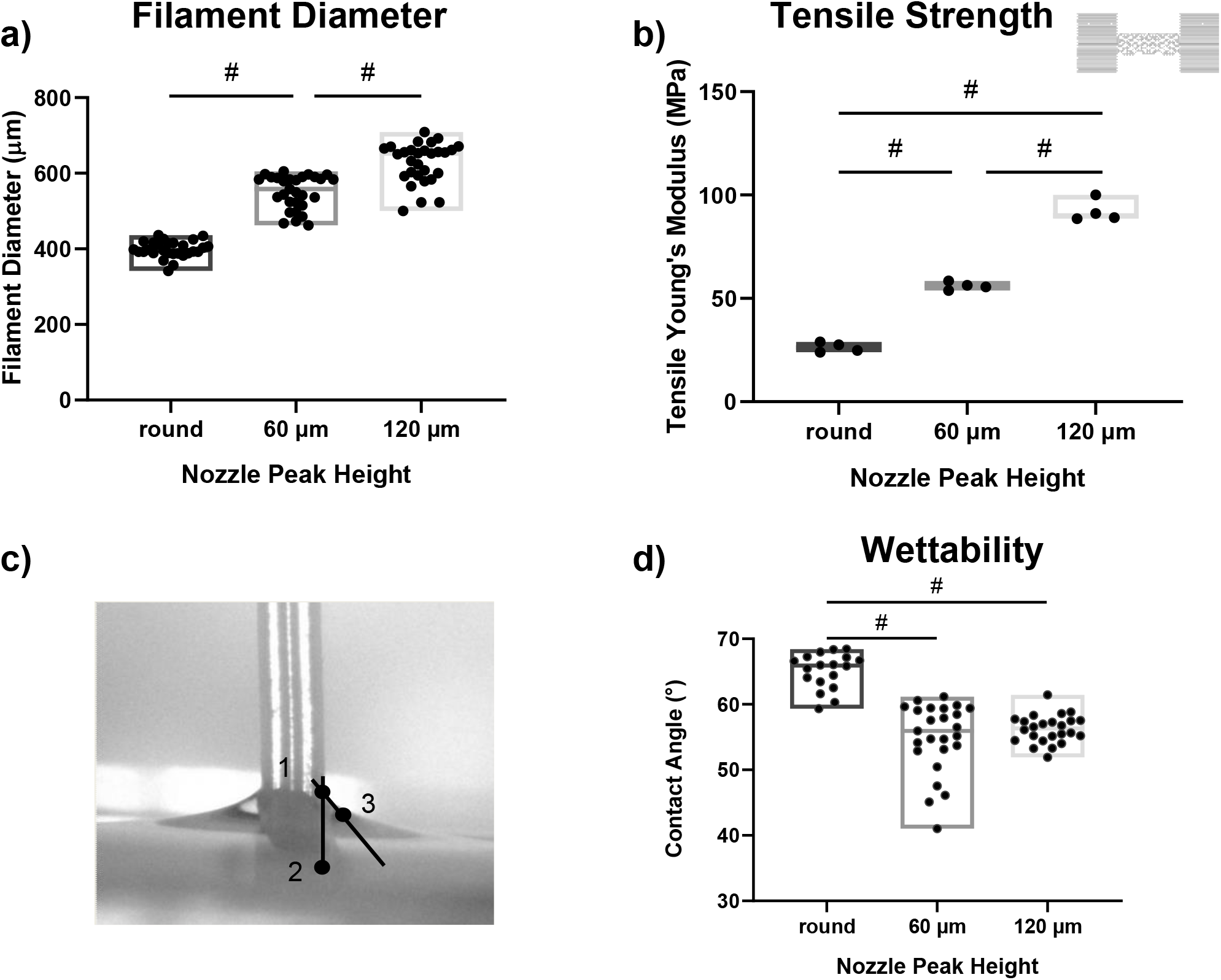
Results of scaffold characterisation show nozzle dependent differences. **(a)** SEM pictures show a increase in filament diameter with topographical features. (# p-value < 0.05) **(b)** Scaffolds printed with patterned nozzles exhibit increased circumferential tensile Young’s moduli. (# p-value < 0.05) **(c)** Contact angle measurement was performed by using three predefined points: 1) contact point between water and material, 2) point on the PCL filament and 3) point at the steepest slope on the water meniscus curvature. **(d)** Contact angle measurements indicated decreases in contact angle, or increases in wettability, with larger topographical features. (# p-value < 0.05).

#### 2.2.4 Wettability

Wettability was determined using contact angle measurement. A typical droplet size in contact angle measurement on PCL ranges from 2-10 μl (59-68), which would cover the whole surface of a 3D printed filament. Therefore, extruded PCL filaments were immersed in Milli-Q^®^ water and pictures were continuously taken while raising the filaments slowly from the water with DinoCapture Camera (AnMo Electronic Corporation, Taipei, Taiwan). Contact angle was measured on the pictures (n ≥ 18) with AxioVision4 (version 4.9.1.0, Carl Zeiss Microscopy, Jena, Germany). Three points were defined to measure contact angle at the water material interface: 1) contact point between water and filament, 2) point on the PCL filament, and 3) point at the steepest slope on the water meniscus curvature (***Figure 2(c)***).

### 2.3 Cell Culture

Bone marrow derived mesenchymal stem cells (BM-MSCs) from three healthy donors (one male (donor 1), age 85, and two females (donor 2 and 3), ages 33 and 55) were isolated and cryopreserved at a density of 2 × 10^6^ cells per ml. General Consent which also covers anonymization of health-related data and biological material was obtained. After thawing (passage 1), cells were suspended in growth medium containing α-MEM, 10 % FBS, 1 % Pen/Strep and 5 ng/ml basic fibroblast growth factor (b-FGF, Fitzgerald Industries, Acton, USA) and were seeded in T300 tissue culture flasks (TPP, Trasadingen, Switzerland). Non-adherent cells were removed after 24 hours and medium was exchanged three times per week. Cells were passaged upon reaching 80 % confluency.

Prior to seeding with cells, 3D printed PCL scaffolds were sterilized in ethylene oxide (EtO), placed in ultra-low attachment 24-well plates and pre-wetted in growth medium for 24 hours. Then, third-passage BM-MSCs were trypsinized (0.5 % Trypsin EDTA in PBS) and suspended in growth medium at a concentration of 5 × 10^5^ cells/ml. A total of 2.5 × 10^5^ cells were seeded per scaffold and allowed to attach for 48 hours prior to removing non-adherent cells and changing to differentiation medium. The samples were cultured in AF differentiation medium containing high glucose DMEM (4.5 μg/ml D-Glucose), sodium pyruvate (0.11 μg/ml) and sodium bicarbonate (3.7 μg/ml) with 1 % Pen/Strep, L-ascorbic acid 2 phosphate (50 μg/ml), 1 % ITS, non-essential amino acids (5 mg/ml), dexamethasone (0.04 ng/ml) and either 0 or 10 ng/ ml recombinant TGF-β_3_ (69) (PreproTech, Rocky Hill, USA). Medium was exchanged three times per week and samples were evaluated after 48 hours (week 0) and 4 weeks of culturing in AF differentiation medium.

### 2.4 Cell Morphology and Alignment

#### 2.4.1 Cytoskeletal and Focal Adhesion Staining

To evaluate cellular morphology and alignment, samples were fixed using paraformaldehyde (4 % in PBS) for 20 minutes at room temperature and washed three times with PBS. Samples were incubated with an actin cytoskeleton and focal adhesion staining kit, containing TRITC-conjugated phalloidin, anti-vinculin and DAPI following the manufacturer’s protocol (Millicore, Sigma). Briefly, cells were permeabilized with 0.1 % Triton X-100 in PBS for 5 minutes and incubated with anti-vinculin at a dilution of 1:200. AlexaFluor488 goat anti-mouse secondary antibody (2 mg/ml, Invitrogen, Thermo Fisher Scientific) was double stained with TRITC-conjugated phalloidin at a dilution of 1:200 and incubated for 60 minutes at room temperature. DAPI staining (1:1000) was performed subsequently and samples were washed with wash buffer (1x PBS containing 0.05 % Tween-20) prior to further analysis.

#### 2.4.2 Image Acquisition

Stained samples were analysed as z-stacks (12.48 μm intervall) using confocal microscope (Airyscan LSM800 Confocal Microscope, Zeiss) with the following channels: DAPI (405 nm), Alexa488 (488 nm), Texas red (561 nm), and ESID-T1 (transmitted light). For further analysis z-stacks of fluorescent channels were merged into a single plane with Zen Software (Zen 2.3 (blue edition), Carl Zeiss Microscopy).

#### 2.4.3 Image Analysis

Cell alignment was analysed using fast Fourier transformation (FFT) and “Oval Profile” (70) plugin in ImageJ as previously described by Tognato *et al*. (71). Briefly, confocal images (n ≥ 9) of merged channels were imported in ImageJ and an unsharp mask with radius 1 and mask 0.65 was applied. FFT was performed to display power spectrum in polar coordinates, following rotation by 90 ° reversing intrinsic rotation resulting from FFT. A circular selection on the power spectrum with radius width/4 was selected subsequently. “Oval Profile” plugin was executed and resulting pixel intensities were normalized for further analysis. Grey value profiles were integrated in the region of interest (direction of printing evaluated by ESID channel ± 10 °) and divided by the total area under the curve (AUC) to semi-quantify average degree of alignment.

### 2.5 ECM Production

After 4 weeks of culture, samples were fixed in 4 % paraformaldehyde as described in *section 2.4.1*. Cells were incubated in blocking solution containing 10 % goat serum in 0.1 % PBS-Tween (0.1 % Tween-20 in PBS) and stained with primary antibody against collagen type I (host mouse, 1:5000 in PBS-Tween), decorin (6D6, host mouse, 37 μg/ml, DSHB (Developmental Studies Hybridoma Bank), Iowa, USA) and fibromodulin (FMOD, host rabbit, 0.5 mg/ml, Thermo Fisher Scientific). Secondary antibody AlexaFluor647 (goat anti-mouse or goat anti-rabbit IgG, 2 ng/ml, Invitrogen, Thermo Fisher Scientific) was used at 1:200 dilution in PBS-Tween. Cell nuclei were counterstained with DAPI (500 μg/ml) at a dilution of 1:500 in PBS-Tween. Control samples were stained without primary antibodies to ensure no nonspecific binding of secondary antibody. Samples were analysed using confocal microscopy with the following channels: Alexa647 (647 nm) and DAPI (405 nm). Images were merged as described in *section 2.4.2*.

### 2.6 Glycosaminoglycan and DNA Content

Cell-seeded scaffolds were collected at weeks 0 and 4 and digested in proteinase K solution (0.5 mg/ml in phosphate buffer, pH 6.5, Roche, Basel, Switzerland) for 48 hours at 56 °C. DNA content was measured by using Hoechst 33258 and absorbance at 360 nm (excitation) and 465 nm (fluorescence emission) was read. Calf thymus DNA (100 μg/ml, Invitrogen, Thermo Fisher Scientific) was used to create the DNA standard. Sulphated glycosaminoglycan (sGAG) content was determined using 1,9-dimethyl-methylene blue (16 ng/ml, pH 3). Absorbance was read at 535 nm with a Micro Plate Reader (Tecan, Maennedorf, Switzerland). sGAG concentration was calculated from a standard curve obtained with chondroitin 4-sulfate sodium salt from bovine trachea (1 mg/ml) and normalized to DNA content.

### 2.7 Gene Expression

RNA was isolated from monolayer (day 0) and week 4 scaffolds using TRI-reagent. Three scaffolds were pooled into groups of three samples each and homogenized with a Mixer Mill (MM400, Retsch GmbH, Haan, Germany). TaqMan™ Reverse Transcription Kit was used to generate cDNA from a total 0.5 μg RNA per sample. Relative gene expression reactions were set up in 10 μl reaction mixes using TaqMan™ MasterMix, relevant human primers (***Supplementary Table 1***), diethyl pyrocarbonate treated water (DEPC-water) and cDNA. Real-time polymerase chain reaction (real-time PCR) was performed on duplicates using QuantStudio 7 Flex (Applied Biosystems, Thermo Fisher Scientific, Waltham, USA). The relative gene expression was identified using 2^-ΔΔCt^ value (72, 73) with RPLP0 as endogenous control and week 0 samples were used for normalization.

### 2.8 Statistical Analysis

Quantitative results were analysed and presented using Graph Pad Prism software (Prism 8.1.0). First, datasets were tested for their normality and lognormality. Second, normally distributed results were statistically evaluated by one-way ANOVA with Tukey’s post-hoc. Unless otherwise stated, non gausian distributed data were analysed using Kruskal Wallis test with Dunn’s post hoc. All values are presented as boxplots with median and p-values < 0.05 were considered as significant.

## 3. Results

### 3.1 Material Characterization of 3D Printed Constructs

#### 3.1.1 Light Microscopy

Custom-designed printer nozzles with periodic circumferential patterns of different peak sizes were used to induce aligned surface topography on the 3D printed constructs. Microscopic imaging of extruded filaments revealed that the spaces between the peaks in the nozzle shape resulted in uniaxially aligned surface grooves (***Figure 1(a)-(b)***). Cross-sectional images demonstrated extruded filaments possessing circumferential peaks with average height of 17.87 ± 5.69 μm and 31.46 ± 8.09 μm for the nozzles with 60 and 120 μm peak height, respectively (***Figure 1(c)***).

#### 3.1.2 Scanning Electron Microscopy

Scanning electron microscopy (***Figure 1(d)(e)***) revealed that all nozzles, independent of their shape, produced filaments with slight aligned textures across the surfaces (single arrows in ***Figure 1(e)***), attributed to inherent stretching of the polymer melt post-deposition due to nozzle translation. Extrusion through the patterned nozzles additionally generated distinct uniaxial aligned grooves (double arrows in ***Figure 1(e)***) along the printed filaments that were 10.87 ± 3.09 μm or 17.77 ± 4.91 μm wide for the nozzle with 60 or 120 μm peak height, respectively. Furthermore, filament diameter was calculated to be 388.09 ± 19.20 μm for the round and 583.57 ± 16.77 μm or 667.49 ± 16.03 μm for the nozzles with 60 or 120 μm peak height, respectively (***Figure 2(a)***).

#### 3.1.3 Mechanical characterization

Tensile testing revealed increased Young’s modulus with nozzle peak height (***Figure 2(b)***). Scaffolds printed with the round nozzle exhibited the lowest tensile modulus (26.30 ± 2.03 MPa). Scaffolds printed with the patterned nozzles revealed significantly higher Young’s moduli, with 56.02 ± 1.63 MPa and 92.17 ± 4.66 MPa for the nozzles with 60 and 120 μm peak height, respectively. Statistical analysis revealed significant differences in scaffold tensile properties between the nozzles (p-value < 0.05).

#### 3.1.4 Wettability

Water contact angle measurements revealed that increasing peak height on the PCL filaments produced decreasing water contact angle, indicating more hydrophilic surfaces (74). The patterned nozzles produced surfaces with significantly lower contact angles (p < 0.05) compared to the round nozzle. The average contact angles were 65.09 ± 2.72 ° for scaffolds extruded with the round nozzle and 54.94 ± 5.31 ° or 56.21 ± 2.13 ° for scaffolds extruded with the 60 μm or 120 μm peak height nozzles (***Figure 2(d)***).

### 3.2 Cell Alignment and Proliferation on the 3D Printed Scaffolds

BM-MSCs cultured on the 3D printed scaffolds were stained with anti-vinculin, phalloidin and DAPI and imaged with confocal microscopy. Shown in ***Figure 3 (a)-(c)*** is a representative image of a cell-seeded scaffold printed with each nozzle, the corresponding grey value profile from FFT, and the average calculated degree of alignment. Semi-quantitative analysis revealed that cells seeded on scaffolds printed with the nozzle with 60 μm peak height exhibited the highest degree of alignment in the printing direction (24.46 ± 3.00 %) at week 0 (***Figure 3(a)***). Cells seeded on scaffolds printed with the round nozzle showed an intermediate degree of alignment of 21.65 ± 2.99 %, whereas cells on scaffolds printed with the nozzle with 120 μm peak height exhibited the lowest degree of alignment (19.22 ± 4.02 %). One-way ANOVA did not reveal statistically significant differences in cell alignment between the three nozzles at week 0 (p > 0.05) (***Figure 3(d)***).

**Figure 3:**
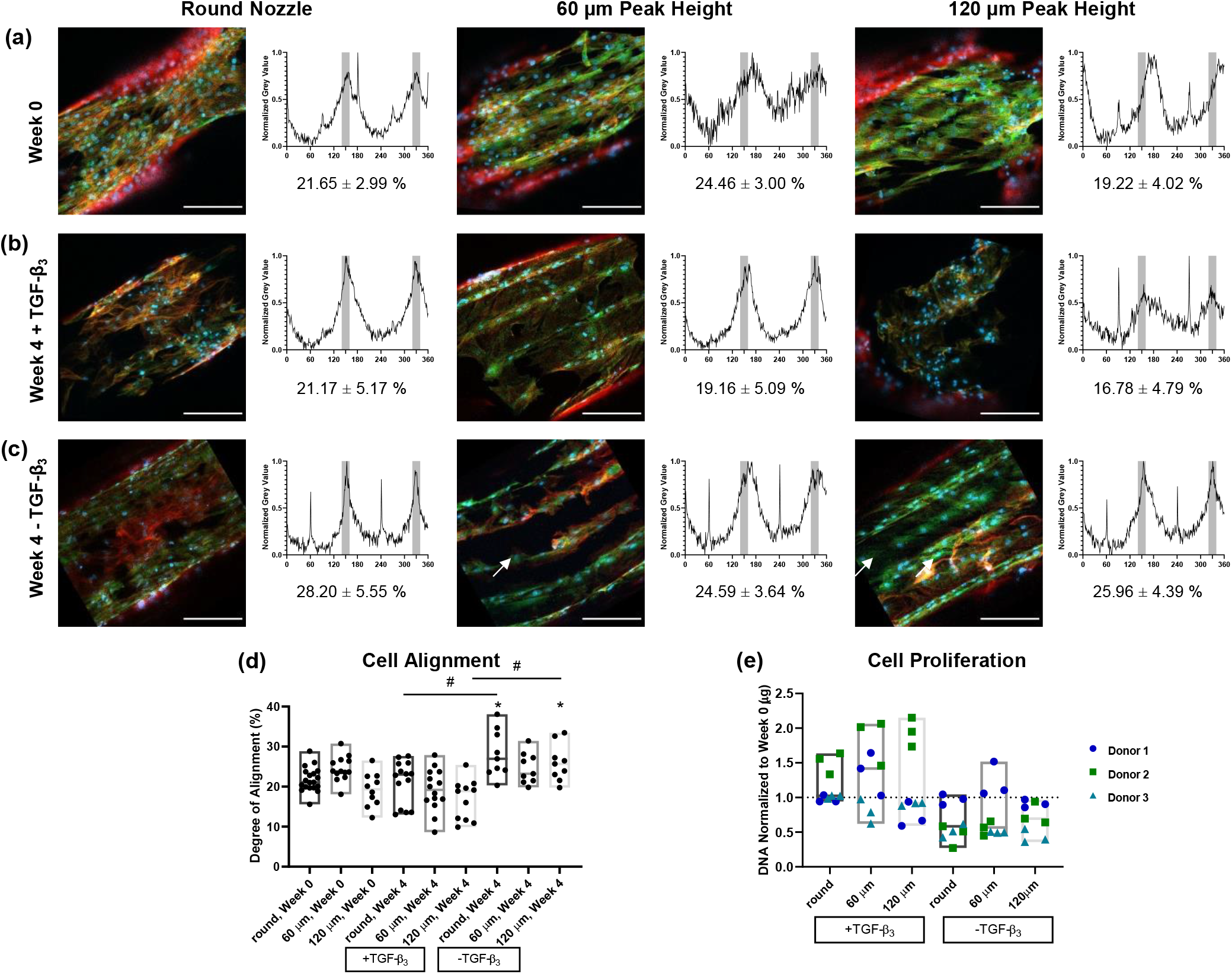
BM-MSCs on PCL scaffolds stained with anti-vinculin (green), phalloidin (red) and DAPI (blue) at week 0 **(a)** and after 4 weeks of culture in medium with **(b)** or without TGF-β_3_ **(c)** with grey value profile resulting from FFT. Degree of alignment was calculated by the portion of AUC at the region of interest (printing direction ± 10 °, highlighted in grey). (Scalebars 200 μm) **(d)** Statistical analysis of degree of alignment revealed no significant differences between nozzles. * indicates p < 0.05 versus week 0, # indicates p < 0.05 between groups. **(e)** DNA content at week 4 normalized to week 0 (donor 1 dark blue circles, donor 2 green squares, donor 3 cyan triangles).

After 4 weeks of culturing in differentiation medium containing growth factor (+TGF-β_3_), cells seeded on scaffolds printed with the round nozzle exhibited the highest degree of alignment (21.17 ± 5.17 %) (***Figure 3(b)***). The cells seeded on the scaffolds printed with the patterned nozzles showed a lower degree of alignment, with 19.16 ± 5.09 % or 16.78 ± 4.79 % for the nozzles with 60 or 120 μm peak height, respectively. After 4 weeks of culture, no significant differences in cell alignment between the round and patterned nozzles (p > 0.05) was observed in the +TGF-β_3_ group (***Figure 3(d)***). Additionally, cells cultured with +TGF-β_3_ did not differ statistically in their degree of alignment at week 4 compared to week 0.

After 4 weeks of culturing in medium without growth factor (-TGF-β_3_) (***Figure 3(c)***), cells seeded on scaffolds printed with the round nozzle induced the highest degree of cellular alignment (28.20 ± 5.55 %). Cells on scaffolds printed with nozzles possessing 60 and 120 μm peak height showed slightly lower degrees of alignment compared to the round nozzle (24.59 ± 3.64 % and 25.96 ± 4.39 %, respectively, p > 0.05). Cells in the -TGF-β_3_ group on scaffolds printed with the round and 120 μm peak height nozzles increased their degree of alignment compared to week 0 (p < 0.05). Also at week 4 of culture, cells on scaffolds printed with the round and 120 μm peak height nozzles exhibited significantly higher degrees of alignment cultured without TGF-β_3_ compared to with TGF-β_3_ (p < 0.05, ***Figure 3(d)***).

Wang and colleagues analysed cellular alignment over 180 ° and concluded that a degree of cellular alignment greater than 11.7 % is defined as anisotropic (75). In an isotropic material, cells are equally distributed and 0.56 % of the cells will be located in a region of 1 °. By applying this concept into our area of interest (direction of printing ± 10 °) a degree of alignment above 23.40 % indicates anisotropic alignment. Thus, it can be determined that anisotropic alignment was observed on scaffolds printed with all of the nozzles when cultured in the absence of TGF-β_3_. Differences in cellular alignment between the +TGF-β_3_ and -TGF-β_3_ groups may be explained within the context of cell proliferation. Compared to week 0, scaffolds cultured in the absence of TGF-ß_3_ showed a decreasing trend in DNA content (p > 0.05), whereas scaffolds cultured in the presence of TGF-ß_3_ exhibited an increasing trend (p > 0.05) (***Figure 3(e)***). These results point towards a necessity for controlling seeding density when using topography to modulate patterning.

When patterning was conserved, the round nozzle topography induced a continuous, web-like expression of actin and vinculin across the printed surfaces. On the filaments printed with patterned nozzles, the cells had an intrinsic tendency to settle within the concave grooves, grouped end-to-end within longitudinal rows that were one (60 μm peak height nozzle) or two (120 μm peak height nozzle) cells wide (white arrows in ***Figure 3(c)***).

### 3.3 ECM Production

Confocal immunofluorescence was used for qualitative evaluation of extracellular matrix production on the scaffolds at 4 weeks of culture. Deposition of collagen I, the major extracellular matrix component of the outer AF, and small leucine rich proteoglycans (SLRPs), decorin and fibromodulin, corresponded with cellular organization on the scaffolds (***Figure 4 (a)***). Again, patterning was better conserved in the absence of TGF-β_3_. The presence of sGAG on the scaffolds was verified quantitatively with a 1,9-dimethyl-methylene blue assay. Total sGAG, when normalized to DNA content on the scaffolds, increased among all nozzles compared to week 0, however one-way ANOVA revealed that the increase was statistically significant only for the +TGF-β_3_ group (p < 0.05). Additionally, the +TGF-β_3_ group produced significantly more (p < 0.05) sGAG compared to the -TGF-β_3_ group (***Figure 4 (b)***) for all the nozzle patterns.

**Figure 4:**
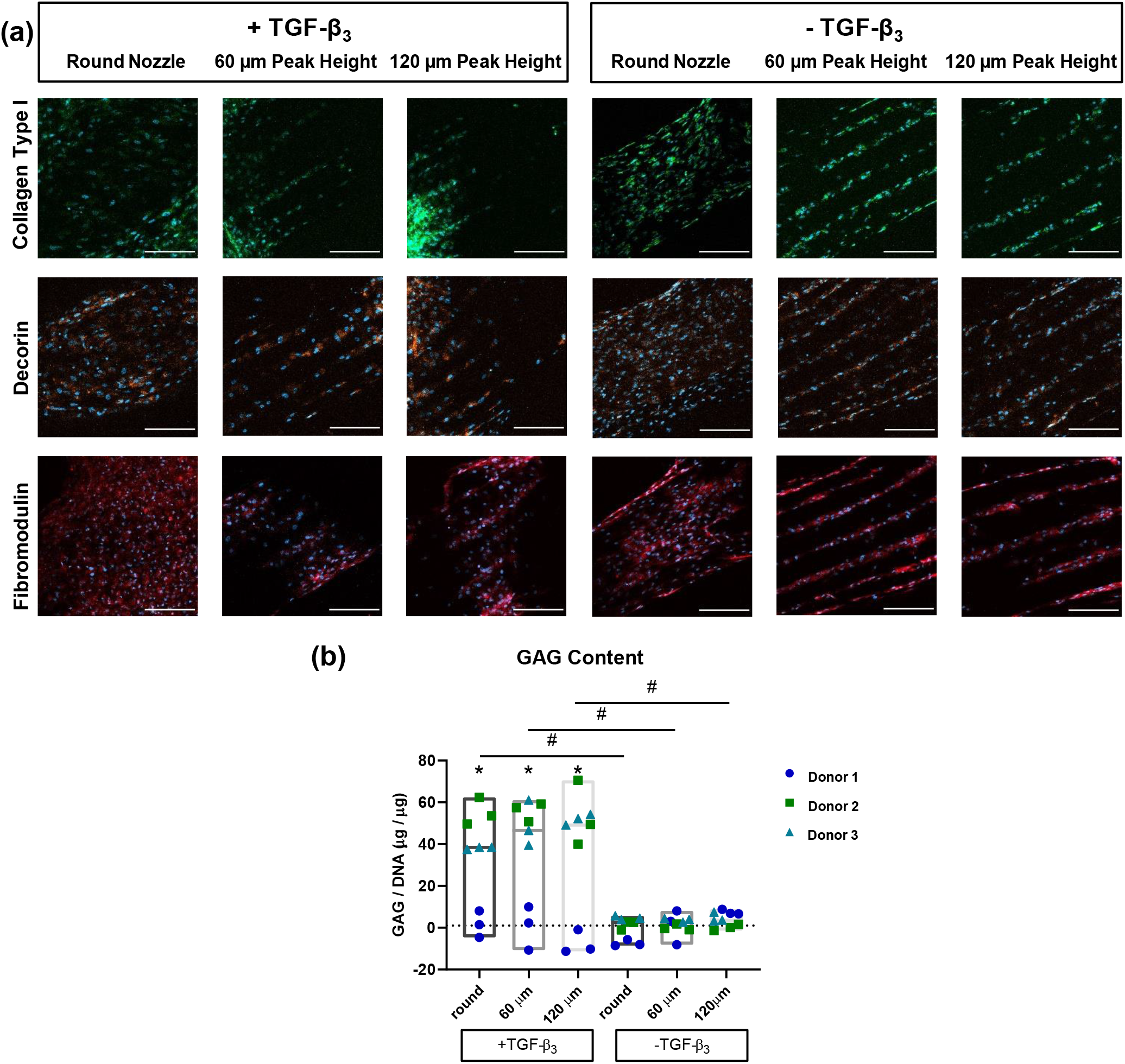
**(a)** Representative pictures of collagen type I (green), decorin (orange), fibromodulin (red) and DAPI (blue) for BM-MSCs (all donors) on PCL scaffolds after 4 weeks of culture in medium with or without TGF-ß_3_. Deposition of ECM components closely followed patterns observed for cellular organization. (Scalebars 200 μm) **(b)** Quantitative measurement of sGAG per DNA on the scaffolds indicated higher sGAG production for the group cultured with TGF-β_3_ in the medium. (donor 1 dark blue circles, donor 2 green squares, donor 3 cyan triangles) * indicates p < 0.05 versus week 0, # indicates p < 0.05 between groups.

### 3.4 Gene Expression

The gene expression study evaluated markers specific to the outer AF: collagen type I (COL1) (76), SFRP2 (76), collagen type XII (COL12) (76, 77), Mohawk (MKX) (78), CD146 (MCAM) (79), scleraxis (SCX) (80), and transgelin (TAGLN) (79) and markers associated with fibrillogenesis: fibromodulin (FMOD) and decorin (DCN) (77) (***Figure 5***). Trending upregulation was observed for all markers compared to week 0 regardless of TGF-β_3_ supplementation, with statistically significant increases (p < 0.05) for AF specific markers SFRP2 and MKX, in the +TGF-β_3_ cells cultured on scaffolds printed with the round and 60 μm nozzle peak heights. DCN followed the same trend with statistically significant increases in its expression compared to day 0 for the +TGF-β_3_ scaffolds printed with the round and 60 μm nozzle peak heights. FMOD was significantly upregulated compared to week 0 for all the nozzles in the presence of TGF-β_3_ (p < 0.05). FMOD was the only marker exhibiting a significantly higher upregulation for the +TGF-β_3_ compared to -TGF-β_3_ group. TAGLN expression was higher for the -TGF-β_3_ group on scaffolds printed with the patterned nozzles compared to the +TGF-β_3_ group on scaffolds printed with the round nozzle (p < 0.05). COL1 and SCX exhibited a trend with nozzle pattern, with increasing expression with decreasing size of the topography, yet the differences were not statistically significant (p > 0.05).

**Figure 5:**
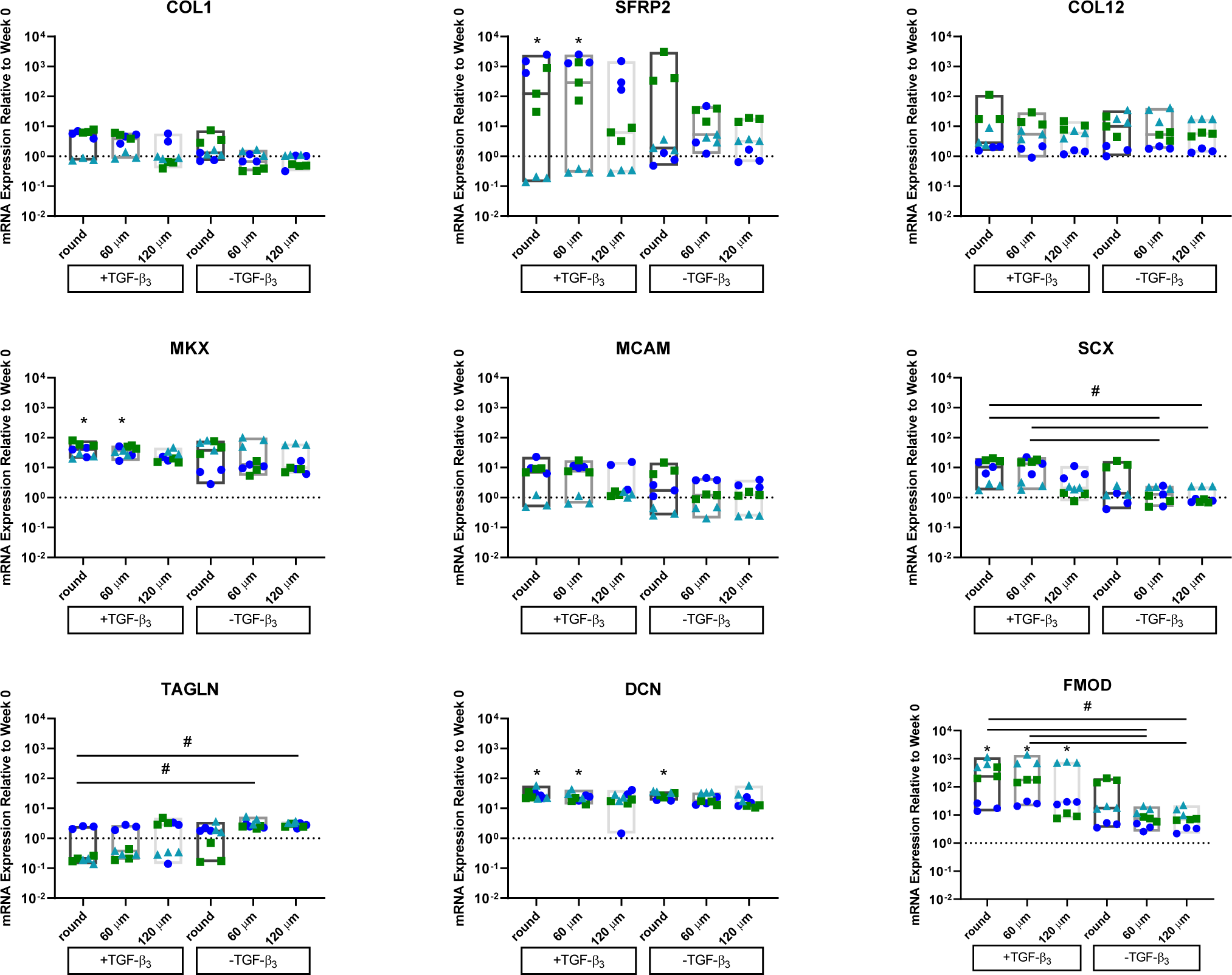
Gene expression data after 4 weeks of culture, normalized to the endogenous control and expression level of cells before starting scaffold culture (week 0). (Donor 1 dark blue circles, Donor 2 green squares, Donor 3 cyan triangles) * indicates p < 0.05 versus week 0, # indicates p < 0.05 between groups.

## 4. Discussion

Low back pain is the most common cause of disability world-wide and closely associated to IVD degeneration (2, 3). Partially to blame for the inadequacies in the current treatment options is the lack of realistic 3D *in vitro* models that recreate native AF architecture. The morphological organization of AF tissue is established during development and its process of maturation is similar to other connective tissues like tendon (34). In the embryo, collagen fibrils initially fuse within membrane bound compartments of a single cell and grow longitudinally as they coalesce within compartments formed between neighboring cells that have aligned nuclei and are stacked end-to-end (55, 57, 81). In fact, tissue morphology is determined by the number and spatial arrangement of cell-to-cell contacts during the developmental phase (57). Within this context, a limitation of current biomaterial-based tissue engineering strategies for the AF is the lack of control over end-to-end and nuclear alignment of seeded cells.

Extrusion-based 3D printing is a widely-adopted, cost-effective, and translational method of scaffold fabrication. By extruding molten PCL through nozzles possessing sinusoidal circumferential geometries, we fabricated 3D angle-ply architectures possessing uniaxial microgrooves that cued the end-to-end longitudinal alignment of cells along the printed filaments. More specifically, extrusion through nozzles with 60 or 120 μm peak height produced grooves that were approximately 10 or 17 μm wide, respectively. Seeded BM-MSCs spontaneously grouped end-to-end in rows of one or a few cells within the microgrooves, an effect that was most pronounced in the -TGF-β_3_ group where cell density was lowest. This longitudinal grouping behavior contrasted with observations for the round (control) nozzle, where a slight aligned texture on the PCL surface induced a continuous web of elongated cells without end-to-end alignment.

The immunofluorescent staining showed aligned phalloidin and vinculin localized into columnar patterns within the microgrooves on scaffolds printed with the 60 or 120 μm nozzle peak heights. Vinculin is a key mechanosensitive protein within cell-matrix and cell-cell adhesions (82), whose function is necessary for the generation of tensional forces across a cell and the assembly of aligned actin (83). Overall, the immunofluorescent images point towards the presence of cytoskeletal tensional forces across cells aligned within the microgrooves. Interestingly, gene expression for the actin-filament associated protein transgelin (TAGLN), which is linked to Myosin II motor generated cytoskeletal tension (84), was highest for scaffolds with microgrooves and cultured in the absence of TGF-β_3_, indicating that end-to-end alignment may favor its upregulation. We speculate that the microgrooves serve as biophysical cues that can be leveraged to amplify tensional forces across the long axis of end-to-end cells. This in turn could drive higher cytoskeletal anisotropy and improve collagen organization compared to a web-like pattern, where tensile forces are shared laterally with many neighbors.

Regarding ECM formation, type I collagen, and the SLRPs, decorin and fibromodulin, were detected on the scaffolds, indicating potential to support AF fiber assembly. It is likely that medium composition can be optimized to further upregulate AF markers and ECM expression. Besides soluble factors, another important consideration affecting ECM is the geometry of the microgrooves. Prior studies indicate that adherent cells position themselves within concave valleys (85). This is consistent with our observation that the BM-MSCs had a tendency to group within the concave microgrooves generated by the patterned nozzles. Essentially then, the microgrooves are the primary sites for collagen nucleation on the scaffolds. The radius of curvature of the grooves can affect cytoskeletal adaptation, gene expression, and ECM morphology (86). Groove geometry should be strongly considered in future studies, not only within the context of downstream matrix assembly, but also when making comparisons between scaffolds printed with different nozzles. While the current data does not allow an elucidation of which geometry is optimal for AF tissue repair, nozzle features (round versus square, peak height, and peak shape) can be tailored to potentially drive a targeted cell behavior.

The biofabrication method presented herein is versatile in terms of the biophysical cues offered to the cells. Scaffold mechanical properties can have a myriad of implications for engineered tissue organization. The 3D printed scaffolds in the current study exhibited tensile moduli in the range of 26.30 ± 2.03 MPa to 92.17 ± 4.66 MPa, which is similar to that of a single outer AF lamella (13.2 ± 5.0 to 82 ± 43 MPa (87-90). While high bulk mechanical properties can be favorable for load-bearing tissue replacement, the deformability of a cell’s underlying matrix is necessary for collagen crimp formation (91). This method is likely adaptable for use with other extrudable polymer systems. PCL has been combined with more elastic components like poly(glycerol sebacate) (92) or collagen (93), resulting in scaffolds that can potentially accommodate a greater degree of cell contraction than PCL alone. Soft hydrogels could potentially be printed with this method, provided there is a mechanism for attaining efficient shape fidelity upon nozzle extrusion. We speculate that the end-to-end patterning of cells on materials possessing a greater degree of deformability could be a potent strategy for synergistically improving fibrillar tissue hierarchical structure and functionality.

Still, there are limitations with the proposed method for building 3D model systems. The topography is dependent on rate of solidification post-deposition; thus, printing results are prone to variations if environmental conditions, such as temperature, fluctuate. While nozzle shape can be modified to produce any continuous feature along an extruded surface, geometric shapes like islands (94), finger-like projections (95), or holes (96) cannot be produced from this single-step extrusion method. Despite these considerations, we propose the biofabrication method can serve as a viable study platform that may lead to the improvement and/or acceleration of aligned collagen fiber assembly for AF repair.

## 5. Conclusion

We described a novel biofabrication method where PCL was extruded through nozzles possessing circumferential sinusoidal peaks, producing multilayered, angle-ply scaffolds with uniaxial microscale grooves. The influence of the grooved topography on BM-MSC alignment, differentiation and ECM expression was investigated. It was demonstrated that BM-MSCs tended to group into longitudinal rows within the grooves, mimicking the end-to-end arrangement of cells during embryonic fibrillogenesis. In contrast, extrusion of PCL through a round control nozzle produced filaments with slight aligned surface textures inducing a continuous web of elongated cells without end-to-end alignment. Collagen I, decorin and fibromodulin were detected on the scaffolds in patterns following cellular organization. Taken together, we present a single-step method for biofabricating scaffolds that spontaneously induce longitudinal cellular alignment in 3D space. We present this work as a potential tool for investigating mechanoregulatory aspects of AF repair, with potential wider relevance to other tissues with aligned microarchitectures.

## 6. Funding acknowledgement

This work was supported by an ON/EORS Kick-Starter Grant (project number 19-185) and AO Spine; NK and GUA were supported by an AO Foundation Fellowship from ETH Zürich Foundation.

## Supplementary Information for Materials and Methods

### Fourier Transform Infrared Spectroscopy

PCL filaments extruded through nozzles with different peak heights were analysed by Fourier transformed infrared spectroscopy (FTIR). The measurements were performed with an attenuated total reflectance-FTIR spectroscope (Burge IR Tensor 27 Version 6.5 Build: 6.5.97) at a wavelength of 400-4000 cm^-1^. Characteristic peaks were marked and area under the curve was calculated with OPUS Spectroscopic software (Version 6.5 Build: 6.5.97, Bruker Optik GmbH, Ettlingen, Germany).

### Gene Expression Analysis

**Supplementary Table 1:**
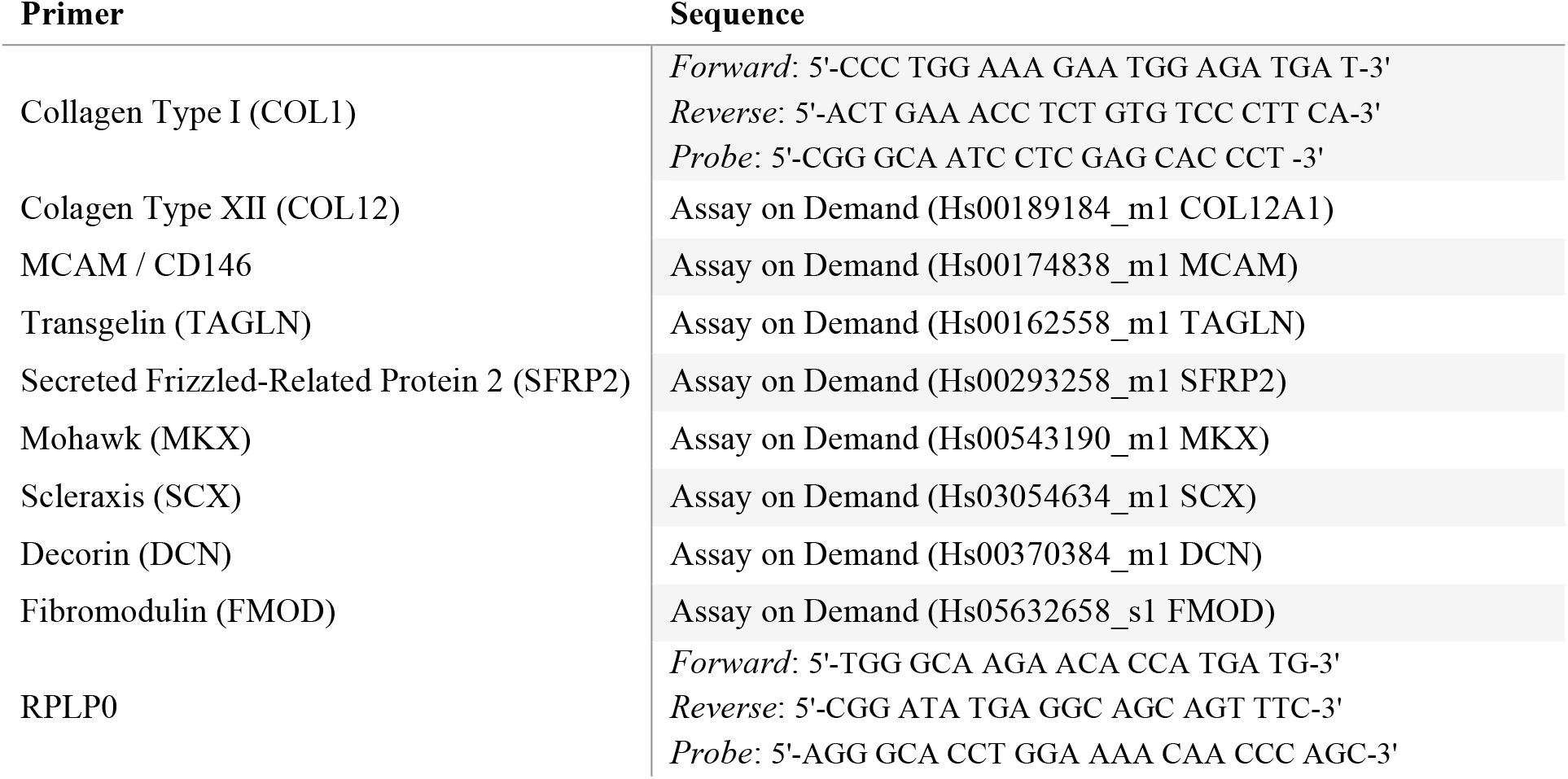
Human Primer Sequences for real-time PCR

## Supplementary Figures

**Supplementary Figure 1:**
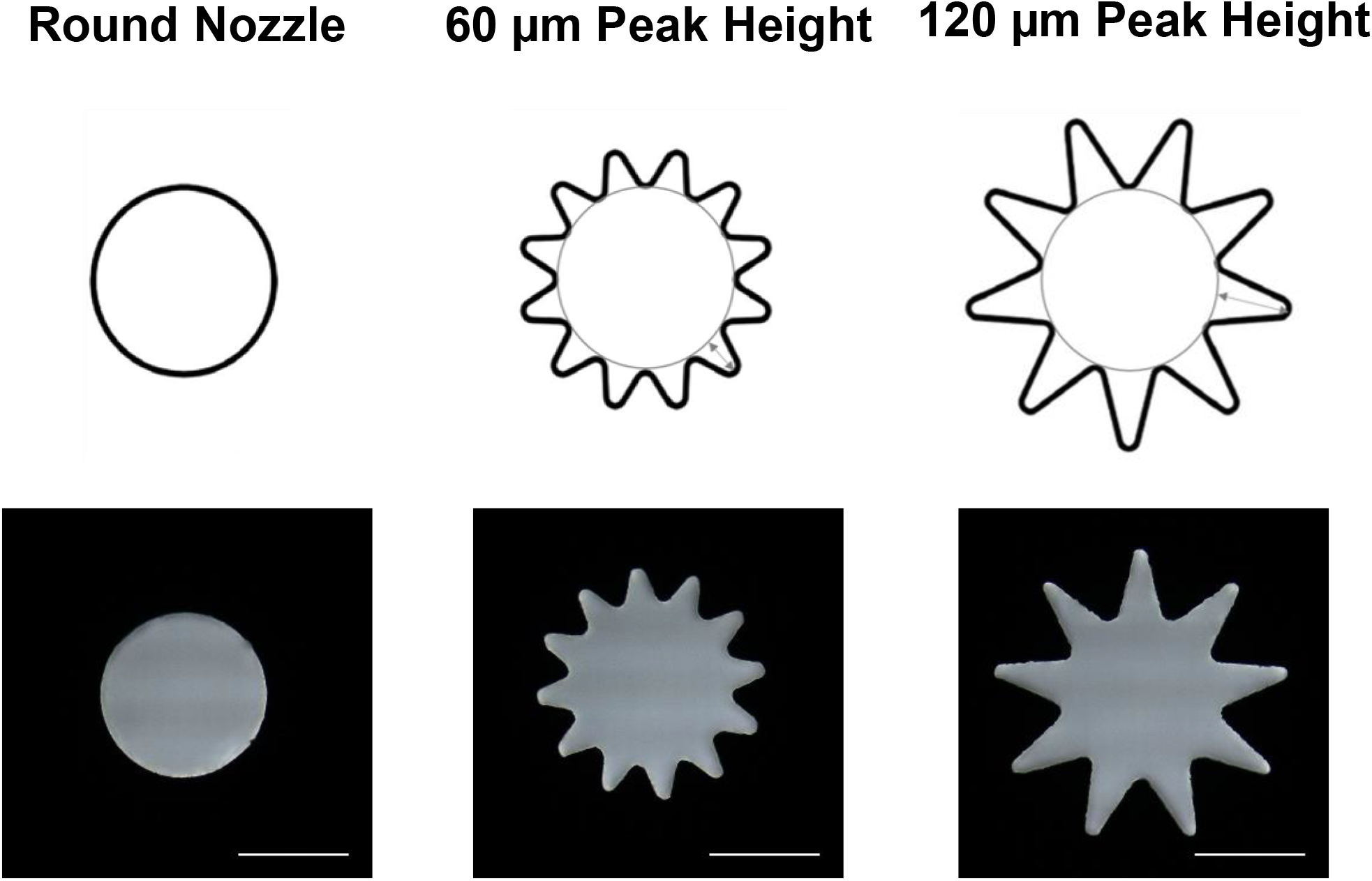
**(a)** Schematics of nozzle geometry and (b) light microscopy images of the engineered 3D printing nozzles with inner diameter of 300 μm and circumferential sinusoidal patterns with 60 and 120 μm peak height (Scalebars 200 μm).

**Supplementary Figure 2:**
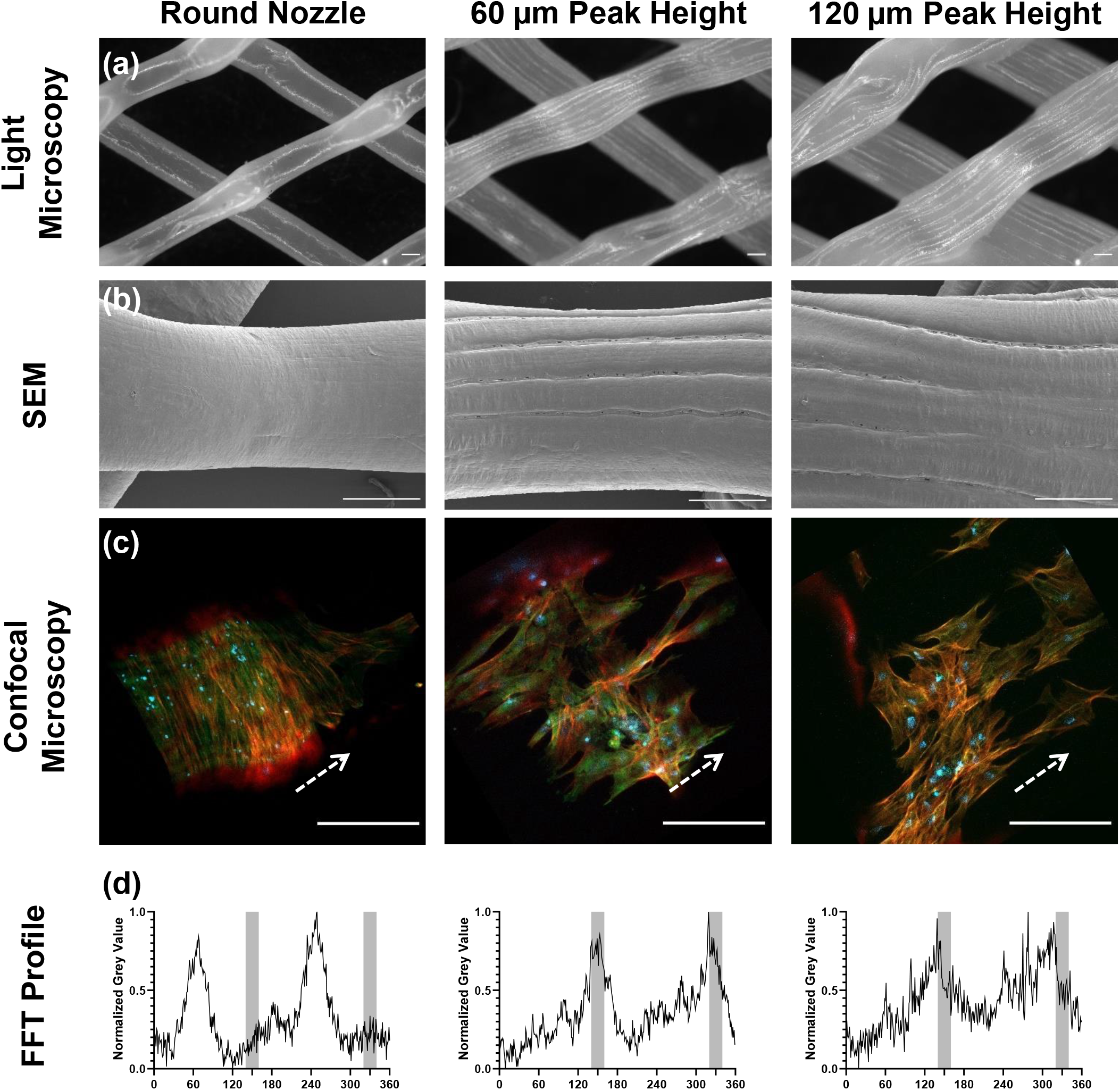
Printing with 8 mm/s resulted in sagging of the filaments interfering topographical guidance. **(a)** Light microscopy top-view images of the uniaxial aligned surface topographies printed with 8 mm/s. **(b)** Scanning electron microscopy top-view images of the surface topographies. **(c)** MSCs on PCL scaffolds stained with anti-vinculin (green), phalloidin (red) and DAPI (blue) after 4 weeks culturing in medium with TGF-β_3_. The white arrows indicate the direction of printing. **(d)** Grey value profile resulting from FFT. (Scalebars 200 μm)

**Supplementary Figure 3:**
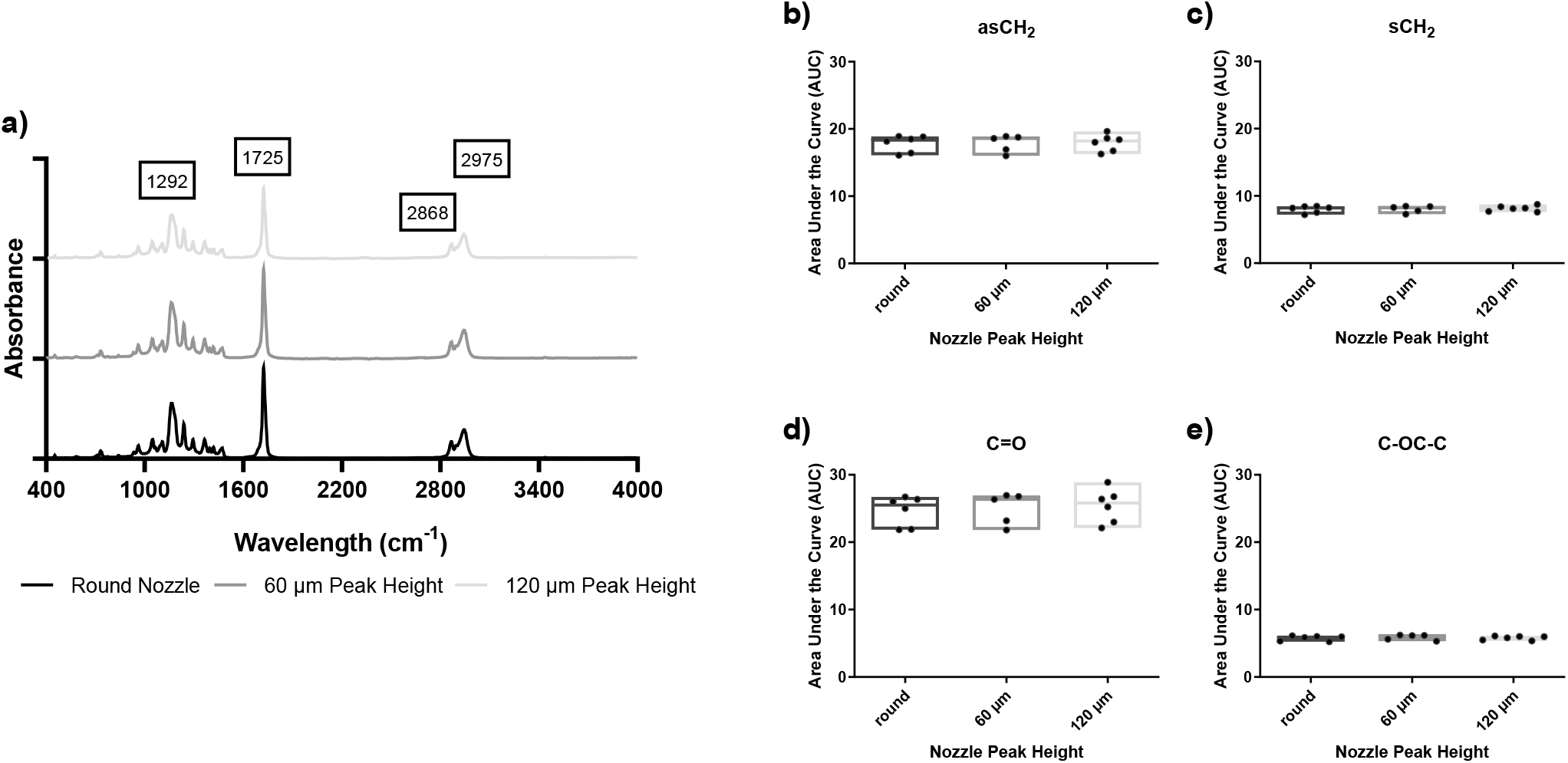
Fourier-transform infrared spectroscopy (FTIR) of PCL filaments revealed characteristic signal peaks. The typical signal peaks with their wavelengths are highlighted **(a)**. FTIR indicated no differences in the area under the curve as a function of nozzle geometry for characteristical bands: asymmetric **(b)** and symmetric **(c)** CH_2_ (2975 and 2868 cm^-1^), C-O backbone (1292 cm^-1^) **(d)** and carbonyl stretching (1725 cm^-1^) **(e)**.

## References

1. Cieza A, Causey K, Kamenov K, Hanson SW, Chatterji S, Vos T. Global estimates of the need for rehabilitation based on the Global Burden of Disease study 2019: a systematic analysis for the Global Burden of Disease Study 2019. The Lancet. 2020;396(20167):2006–17.

2. Wu A, March L, Zheng X, Huang J, Wang X, Zhao J, et al. Global low back pain prevalence and years lived with disability from 1990 to 2017: estimates from the Global Burden of Disease Study 2017. Annals of translational medicine. 2020;8(6):299.

3. Vos T, Barber RM, Bell B, Bertozzi-Villa A, Biryukov S, Bolliger I, et al. Global, regional, and national incidence, prevalence, and years lived with disability for 301 acute and chronic diseases and injuries in 188 countries, 1990–2013: a systematic analysis for the Global Burden of Disease Study 2013. The lancet. 2015;386(9995):743–800.

4. Ito K, Creemers L. Mechanisms of intervertebral disk degeneration/injury and pain: a review. Global Spine J. 2013;3(3):145–52.

5. Erwin WM. Biologically based therapy for the intervertebral disk: who is the patient? Global Spine J. 2013;3(3):193–200.

6. Cheung KM, Karppinen J, Chan D, Ho DW, Song YQ, Sham P, et al. Prevalence and pattern of lumbar magnetic resonance imaging changes in a population study of one thousand forty-three individuals. Spine (Phila Pa 1976). 2009;34(9):934–40.

7. Chan SC, Gantenbein-Ritter B. Intervertebral disc regeneration or repair with biomaterials and stem cell therapy--feasible or fiction? Swiss Med Wkly. 2012;142:w13598.

8. Pattappa G, Li Z, Peroglio M, Wismer N, Alini M, Grad S. Diversity of intervertebral disc cells: phenotype and function. J Anat. 2012;221(6):480–96.

9. Bogduk N, McGuirk B. Medical management of acute chronic low back pain : an evidence-based approach. Amsterdam ; Boston: Elsevier; 2002. viii, 224 p. p.

10. Yang X, Li X. Nucleus pulposus tissue engineering: a brief review. European Spine Journal. 2009;18(11):1564–72.

11. Roughley PJ. Biology of intervertebral disc aging and degeneration: involvement of the extracellular matrix. Spine. 2004;29(23):2691–9.

12. Gu BK, Choi DJ, Park SJ, Kim Y, Kim C. 3D Bioprinting Technologies for Tissue Engineering Applications. Cutting-Edge Enabling Technologies for Regenerative Medicine Advances in Experimental Medicine and Biology. 1078. Singapore2018.

13. Cauble MA, Mancini NS, Kalinowski J, Lykotrafitis G, Moss IL. Atomic force microscopy imaging for nanoscale and microscale assessments of extracellular matrix in intervertebral disc and degeneration. JOR spine. 2020;3(3):e1125.

14. Kos N, Gradisnik L, Velnar T. A brief review of the degenerative intervertebral disc disease. Medical Archives. 2019;73(6):421.

15. Iatridis JC, Michalek AJ, Purmessur D, Korecki CL. Localized intervertebral disc injury leads to organ level changes in structure, cellularity, and biosynthesis. Cellular and molecular bioengineering. 2009;2(3):437–47.

16. Choi H, Tessier S, Silagi ES, Kyada R, Yousefi F, Pleshko N, et al. A novel mouse model of intervertebral disc degeneration shows altered cell fate and matrix homeostasis. Matrix Biology. 2018;70:102–22.

17. Parker SL, Mendenhall SK, Godil SS, Sivasubramanian P, Cahill K, Ziewacz J, et al. Incidence of low back pain after lumbar discectomy for herniated disc and its effect on patient-reported outcomes. Clinical Orthopaedics and Related Research. 2015;473(6):1988–99.

18. Yorimitsu E, Chiba K, Toyama Y, Hirabayashi K. Long-term outcomes of standard discectomy for lumbar disc herniation: a follow-up study of more than 10 years. Spine. 2001;26(6):652–7.

19. de Schepper EI, Damen J, van Meurs JB, Ginai AZ, Popham M, Hofman A, et al. The association between lumbar disc degeneration and low back pain: the influence of age, gender, and individual radiographic features. Spine. 2010;35(5):531–6.

20. Jarman JP, Arpinar VE, Baruah D, Klein AP, Maiman DJ, Muftuler LT. Intervertebral disc height loss demonstrates the threshold of major pathological changes during degeneration. European Spine Journal. 2015;24(9):1944–50.

21. Zhou X, Wang J, Huang X, Fang W, Tao Y, Zhao T, et al. Injectable decellularized nucleus pulposus-based cell delivery system for differentiation of adipose-derived stem cells and nucleus pulposus regeneration. Acta biomaterialia,. 2018;81:115–28.

22. Deyo RA, Jarvik JG, Chou R. Low back pain in primary care. BMJ: British Medical Journal 2014;349:g4266.

23. Ren C, Qin R, Li Y, Wang P. Microendoscopic Discectomy Combined with Annular Suture Versus Percutaneous Transforaminal Endoscopic Discectomy for Lumbar Disc Herniation: A Prospective Observational Study. Pain Physician. 2020;23(6):E713–E21.

24. Benzakour A, Benzakour T. Lumbar disc herniation: long-term outcomes after mini-open discectomy. International orthopaedics. 2019;43(4):869–74.

25. Xu G, Zhang C, Zhu K, Bao Z, Zhou P, Li X. Endoscopic removal of nucleus pulposus of intervertebral disc on lumbar intervertebral disc protrusion and the influence on inflammatory factors and immune function. Experimental and therapeutic medicine. 2020;19(1):301–7.

26. Tang S, Rebholz BJ. Does lumbar microdiscectomy affect adjacent segmental disc degeneration? A finite element study. Journal of Surgical Research. 2013;182(1):62–7.

27. Papagiannis GI, Triantafyllou AI, Konstantina YG, Koulouvaris P, Anastasiou A, Papadopoulos EC, et al. Biomechanical factors could affect lumbar disc reherniation after microdiscectomy. Journal of Orthopaedics and Sports Medicine. 2019;1(2):46–50.

28. Pang JY, Tan F, Chen WW, Li CH, Dou SP, Guo JR, et al. Comparison of microendoscopic discectomy and open discectomy for single-segment lumbar disc herniation. World Journal of Clinical Cases. 2020;8:2942.

29. Miller LE, McGirt MJ, Garfin SR, Bono CM. Association of annular defect width after lumbar discectomy with risk of symptom recurrence and reoperation: systematic review and meta-analysis of comparative studies. Spine. 2018;43(5):E308.

30. Miller LE, Allen RT, Duhon B, Radcliff KE. Expert review with meta-analysis of randomized and nonrandomized controlled studies of Barricaid annular closure in patients at high risk for lumbar disc reherniation. Expert review of medical devices. 2020;17(5):461–9.

31. Castillo H, Chintapalli RTV, Boyajian HH, Cruz SA, Morgan VK, Shi LL, et al. Lumbar discectomy is associated with higher rates of lumbar fusion. The Spine Journal. 2019;19(487-492).

32. Yao Y, Liu H, Zhang H, Wang H, Zhang Z, Zheng Y, et al. Risk factors for the recurrent herniation after microendoscopic discectomy. World neurosurgery. 2016;95(451-455).

33. Peredo AP, Gullbrand SE, Smith HE, Mauck RL. Putting the pieces in place: mobilizing cellular players to improve annulus fibrosus repair. Tissue Engineering Part B: Reviews. 2021;27(4):295–312.

34. Cox MK, Serra R. Development of the Intervertebral Disc. In: Shapiro IM, Risbud MV, editors. The Intervertebral Disc: Molecular and Structural Studies of the Disc in Health and Disease Heidelberg, Germany: Springer Verlag Gmbh.; 2016.

35. Hayes AJ, Benjamin M, Ralphs JR. Role of actin stress fibres in the development of the intervertebral disc: cytoskeletal control of extracellular matrix assembly. Developmental dynamics: an official publication of the American Association of Anatomists. 1999;215(3):179–89.

36. Hayes AJ, Isaacs MD, Hughes C, Caterson B, Ralphs JR. Collagen fibrillogenesis in the development of the annulus fibrosus of the intervertebral disc. Eur cell mater. 2011;22:226–41.

37. Chu G, Shi C, Wang H, Zhang W, Yang H, Li B. Strategies for Annulus Fibrosus Regeneration: From Biological Therapies to Tissue Engineering. Front Bioeng Biotechnol. 2018;6:90.

38. Nerurkar NL, Baker BM, Sen S, Wible EE, Elliott DM, Mauck RL. Nanofibrous biologic laminates replicate the form and function of the annulus fibrosus. Nat Mater. 2009;8(12):986–92.

39. Ma J, He Y, Liu X, Chen W, Wang A, Lin CY, et al. A novel electrospun-aligned nanoyarn/three-dimensional porous nanofibrous hybrid scaffold for annulus fibrosus tissue engineering. Int J Nanomedicine. 2018;13:1553–67.

40. Nerurkar NL, Elliott DM, Mauck RL. Mechanics of oriented electrospun nanofibrous scaffolds for annulus fibrosus tissue engineering. J Orthop Res. 2007;25(8):1018–28.

41. Kang R, Svend Le DQ, Li H, Lysdahl H, Chen M, Besenbacher F, et al. Engineered three-dimensional nanofibrous multi-lamellar structure for annulus fibrosus repair. J Mater Chem B. 2013;1(40):5462–8.

42. Shamsah AH, Cartmell SH, Richardson SM, Bosworth LA. Tissue Engineering the Annulus Fibrosus Using 3D Rings of Electrospun PCL:PLLA Angle-Ply Nanofiber Sheets. Front Bioeng Biotechnol. 2019;7:437.

43. Shamsah AH, Cartmell SH, Richardson SM, Bosworth LA. Mimicking the Annulus Fibrosus Using Electrospun Polyester Blended Scaffolds. Nanomaterials (Basel). 2019;9(4).

44. Zhou P, Chu G, Yuan Z, Wang H, Zhang W, Mao Y, et al. Regulation of differentiation of annulus fibrosus-derived stem cells using heterogeneous electrospun fibrous scaffolds. J Orthop Translat. 2021;26:171–80.

45. Liu C, Zhu C, Li J, Zhou P, Chen M, Yang H, et al. The effect of the fibre orientation of electrospun scaffolds on the matrix production of rabbit annulus fibrosus-derived stem cells. Bone Res. 2015;3:15012.

46. Gluais M, Clouet J, Fusellier M, Decante C, Moraru C, Dutilleul M, et al. In vitro and in vivo evaluation of an electrospun-aligned microfibrous implant for Annulus fibrosus repair. Biomaterials. 2019;205:81–93.

47. Wismer N, Grad S, Fortunato G, Ferguson SJ, Alini M, Eglin D. Biodegradable electrospun scaffolds for annulus fibrosus tissue engineering: effect of scaffold structure and composition on annulus fibrosus cells in vitro. Tissue Eng Part A. 2014;20(3-4):672-82.

48. Duan G, Jiang S, Jérôme V, Wendorff JH, Fathi A, Uhm J, et al. Ultralight, soft polymer sponges by self-assembly of short electrospun fibers in colloidal dispersions. Advanced Functional Materials. 2015;25(19):2850–6.

49. Huang R, Gao X, Wang J, Chen H, Tong C, Tan Y, et al. Triple-layer vascular grafts fabricated by combined E-jet 3D printing and electrospinning. Annals of biomedical engineering. 2018;46(9):1254–66.

50. Gullbrand SE, Kim DH, Bonnevie E, Ashinsky BG, Smith LJ, Elliott DM, et al. Towards the scale up of tissue engineered intervertebral discs for clinical application. Acta biomaterialia. 2018;70:154–64.

51. Puetzer JL, Ma T, Sallent I, Gelmi A, Stevens MM. Driving hierarchical collagen fiber formation for functional tendon, ligament, and meniscus replacement. Biomaterials. 2021;269:120527.

52. Puetzer JL, Bonassar LJ. Physiologically distributed loading patterns drive the formation of zonally organized collagen structures in tissue-engineered meniscus. Tissue Engineering Part A. 2016;22(13-14):907–16.

53. Puetzer JL, Koo E, Bonassar LJ. Induction of fiber alignment and mechanical anisotropy in tissue engineered menisci with mechanical anchoring. Journal of biomechanics. 2015;48(8):1436–43.

54. Bowles RD, Williams RM, Zipfel WR, Bonassar LJ. Self-assembly of aligned tissue-engineered annulus fibrosus and intervertebral disc composite via collagen gel contraction. Tissue Engineering Part A. 2010;16(4):1339–48.

55. Kapacee Z, Richardson SH, Lu Y, Starborg T, Holmes DF, Baar K, et al. Tension is required for fibripositor formation. Matrix Biology. 2008;27(4):371–5.

56. Cai J, Xie X, Li D, Wang L, Jiang J, Mo X, et al. A novel knitted scaffold made of microfiber/nanofiber core–sheath yarns for tendon tissue engineering. Biomaterials Science. 2020;8(16):4413–25.

57. Kalson NS, Lu Y, Taylor SH, Starborg T, Holmes DF, Kadler KE. A structure-based extracellular matrix expansion mechanism of fibrous tissue growth. Elife. 2015;4:e05958.

58. Richardson SH, Starborg T, Lu, Y., Humphries SM, Meadows RS, Kadler KE. Tendon development requires regulation of cell condensation and cell shape via cadherin-11-mediated cell-cell junctions. Molecular and cellular biology. 2007;27(17):6218–28.

59. Hassan MI, Sultana N. Characterization, drug loading and antibacterial activity of nanohydroxyapatite/polycaprolactone (nHA/PCL) electrospun membrane. 3 Biotech. 2017;7(4):249.

60. Kosorn W, Sakulsumbat M, Lertwimol T, Thavornyutikarn B, Uppanan P, Chantaweroad S, et al. Chondrogenic phenotype in responses to poly(varepsilon-caprolactone) scaffolds catalyzed by bioenzymes: effects of surface topography and chemistry. J Mater Sci Mater Med. 2019;30(12):128.

61. Andrade TM, Mello DCR, Elias CMV, Abdala JMA, Silva E, Vasconcellos LMR, et al. In vitro and in vivo evaluation of rotary-jet-spun poly(varepsilon-caprolactone) with high loading of nano-hydroxyapatite. J Mater Sci Mater Med. 2019;30(2):19.

62. Lobo AO, Afewerki S, de Paula MMM, Ghannadian P, Marciano FR, Zhang YS, et al. Electrospun nanofiber blend with improved mechanical and biological performance. Int J Nanomedicine. 2018;13:7891–903.

63. Unal S, Arslan S, Yilmaz BK, Oktar FN, Ficai D, Ficai A, et al. Polycaprolactone/Gelatin/Hyaluronic Acid Electrospun Scaffolds to Mimic Glioblastoma Extracellular Matrix. Materials (Basel). 2020;13(11).

64. Jager A, Donato RK, Perchacz M, Donato KZ, Stary Z, Konefal R, et al. Human metabolite-derived alkylsuccinate/dilinoleate copolymers: from synthesis to application. J Mater Chem B. 2020;8(43):9980–96.

65. Can-Herrera LA, Oliva AI, Dzul-Cervantes MAA, Pacheco-Salazar OF, Cervantes-Uc JM. Morphological and Mechanical Properties of Electrospun Polycaprolactone Scaffolds: Effect of Applied Voltage. Polymers (Basel). 2021;13(4).

66. Costa Salles TH, Volpe-Zanutto F, de Oliveira Sousa IM, Machado D, Zanatta AC, Vilegas W, et al. Electrospun PCL-based nanofibers Arrabidaea chica Verlot -Pterodon pubescens Benth loaded: synergic effect in fibroblast formation. Biomed Mater. 2020;15(6):065001.

67. Rasti Boroojeni F, Mashayekhan S, Abbaszadeh HA, Ansarizadeh M, Khoramgah MS, Rahimi Movaghar V. Bioinspired Nanofiber Scaffold for Differentiating Bone Marrow-Derived Neural Stem Cells to Oligodendrocyte-Like Cells: Design, Fabrication, and Characterization. Int J Nanomedicine. 2020;15:3903–20.

68. Del Angel-Sanchez K, Borbolla-Torres CI, Palacios-Pineda LM, Ulloa-Castillo NA, Elias-Zuniga A. Development, Fabrication, and Characterization of Composite Polycaprolactone Membranes Reinforced with TiO2 Nanoparticles. Polymers (Basel). 2019;11(12).

69. Blanquer SB, Gebraad AW, Miettinen S, Poot AA, Grijpma DW, Haimi SP. Differentiation of adipose stem cells seeded towards annulus fibrosus cells on a designed poly (trimethylene carbonate) scaffold prepared by stereolithography. Journal of tissue engineering and regenerative medicine. 2017;11(10):2752–62.

70. O’Connell B. Oval Profile Plot 2002 [Available from: https://imagej.nih.gov/ij/plugins/oval-profile.html.

71. Tognato R, Armiento AR, Bonfrate V, Levato R, Malda J, Alini M, et al. A Stimuli-Responsive Nanocomposite for 3D Anisotropic Cell-Guidance and Magnetic Soft Robotics. Advanced Functional Materials. 2018;29(9).

72. Schmittgen TD, Livak KJ. Analyzing real-time PCR data by the comparative C(T) method. Nat Protoc. 2008;3(6):1101–8.

73. Yuan JS, Reed A, Chen F, Stewart CN, Jr. Statistical analysis of real-time PCR data. BMC Bioinformatics. 2006;7:85.

74. Wang QJ, Chung Y-W. Encyclopedia of Tribology2013.

75. Wang ZY, Teo EY, Chong MS, Zhang QY, Lim J, Zhang ZY, et al. Biomimetic three-dimensional anisotropic geometries by uniaxial stretch of poly(epsilon-caprolactone) films for mesenchymal stem cell proliferation, alignment, and myogenic differentiation. Tissue Eng Part C Methods. 2013;19(7):538–49.

76. van den Akker GGH, Koenders MI, van de Loo FAJ, van Lent P, Blaney Davidson E, van der Kraan PM. Transcriptional profiling distinguishes inner and outer annulus fibrosus from nucleus pulposus in the bovine intervertebral disc. Eur Spine J. 2017;26(8):2053–62.

77. Hayes AJ, Isaacs MD, Hughes C, Caterson B, Ralphs JR. Collagen fibrillogenesis in the development of the annulus fibrosus of the intervertebral disc. Eur Cell Mater. 2011;22:226–41.

78. Nakamichi R, Ito Y, Inui M, Onizuka N, Kayama T, Kataoka K, et al. Mohawk promotes the maintenance and regeneration of the outer annulus fibrosus of intervertebral discs. Nat Commun. 2016;7:12503.

79. Nakai T, Sakai D, Nakamura Y, Nukaga T, Grad S, Li Z, et al. CD146 defines commitment of cultured annulus fibrosus cells to express a contractile phenotype. J Orthop Res. 2016;34(8):1361–72.

80. Elsaadany M, Winters K, Adams S, Stasuk A, Ayan H, Yildirim-Ayan E. Equiaxial Strain Modulates Adipose-derived Stem Cell Differentiation within 3D Biphasic Scaffolds towards Annulus Fibrosus. Sci Rep. 2017;7(1):12868.

81. Richardson SH, Starborg T, Lu Y, Humphries SM, Meadows RS, Kadler KE. Tendon development requires regulation of cell condensation and cell shape via cadherin-11-mediated cell-cell junctions. Molecular and cellular biology. 2007;27(17):6218–28.

82. Leerberg JM. Vinculin, cadherin mechanotransduction and homeostasis of cell–cell junctions. Protoplasma. 2013;250(4):817–29.

83. Gilchrist CL, Leddy HA, Kaye L, Case ND, Rothenberg KE, Little D, et al. TRPV4-mediated calcium signaling in mesenchymal stem cells regulates aligned collagen matrix formation and vinculin tension. Proceedings of the National Academy of Sciences. 2019;116(6):1992–7.

84. Liu R, Hossain MM, Chen X, Jin JP. Mechanoregulation of SM22α/transgelin. Biochemistry. 2017;56(41):526–5538.

85. Pieuchot L, Marteau J, Guignandon A, Dos Santos T, Brigaud I, Chauvy PF, et al. Curvotaxis directs cell migration through cell-scale curvature landscapes. Nature communications. 2018;9(1):1–13.

86. Werner M, Blanquer SB, Haimi SP, Korus G, Dunlop JW, Duda GN, et al. Surface curvature differentially regulates stem cell migration and differentiation via altered attachment morphology and nuclear deformation. Advance science. 2017;4(2):1600347.

87. Long RG, Torre OM, Hom WW, Assael DJ, Iatridis JC. Design Requirements for Annulus Fibrosus Repair: Review of Forces, Displacements, and Material Properties of the Intervertebral Disk and a Summary of Candidate Hydrogels for Repair. J Biomech Eng. 2016;138(2):021007.

88. Wagner DR, Lotz JC. Theoretical model and experimental results for the nonlinear elastic behavior of human annulus fibrosus. J Orthop Res. 2004;22(4):901–9.

89. Holzapfel GA, Schulze-Bauer CA, Feigl G, Regitnig P. Single lamellar mechanics of the human lumbar anulus fibrosus. Biomech Model Mechanobiol. 2005;3(3):125–40.

90. Skaggs DL, Weidenbaum M, Iatridis JC, Ratcliffe A, Mow VC. Regional variation in tensile properties and biochemical composition of the human lumbar anulus fibrosus. Spine (Phila Pa 1976). 1994;19(12):1310–9.

91. Herchenhan A, Kalson NS, Holmes DF, Hill P, Kadler KE, Margetts L. Tenocyte contraction induces crimp formation in tendon-like tissue. Biomechanics and modeling in mechanobiology. 2012;11(3):449–59.

92. Yang Y, Lei D, Huang S, Yang Q, Song B, Guo Y, et al. Elastic 3D-printed hybrid polymeric scaffold improves cardiac remodeling after myocardial infarction. Advanced healthcare materials. 2019;8(10):1900065.

93. San Choi J, Lee SJ, Christ GJ, Atala A, Yoo JJ. The influence of electrospun aligned poly (ε-caprolactone)/collagen nanofiber meshes on the formation of self-aligned skeletal muscle myotubes. Biomaterials. 2008;29(19):2899–906.

94. Dalby MJ, Childs S, Riehle MO, Johnstone HJH, Affrossman S, Curtis ASG. Fibroblast reaction to island topography: changes in cytoskeleton and morphology with time. Biomaterials. 2003;24(6):927–35.

95. Altay G, Tosi S, García-Díaz M, Martínez E. Imaging the cell morphological response to 3D topography and curvature in engineered intestinal tissues. Frontiers in bioengineering and biotechnology. 2020;8:294.

96. Karuri NW, Porri TJ, Albrecht RM, Murphy CJ, Nealey PF. Nano-and Microscale Holes Modulate Cell-Substrate Adhesion, Cytoskeletal Organization, and -beta1 Integrin Localization in Sv40 Human Corneal Epithelial Cells. IEEE transactions on nanobioscience,. 2006;5(4):273–80.

